# The impact of telomere shortening on human hippocampal neurogenesis: Implications for cognitive function and psychiatric disorder risk

**DOI:** 10.1101/2020.04.19.049411

**Authors:** Alish B. Palmos, Rodrigo R. R. Duarte, Demelza M. Smeeth, Erin C. Hedges, Douglas F. Nixon, Sandrine Thuret, Timothy R. Powell

## Abstract

Telomere shortening is one hallmark of cell ageing that can limit the proliferative capacity of cell populations and increase risk for age-related disease. It has been hypothesized that short telomeres, and subsequently a limited proliferative capacity of hippocampal progenitor cells, could contribute to smaller hippocampal volumes and impaired cognition, amongst psychiatric disorder patients. The current study employed a systematic, multidisciplinary approach which aimed to model the effects of telomere shortening on human hippocampal neurogenesis, and to explore its relationship with cognition and psychiatric disorder risk. We modelled telomere shortening in human hippocampal progenitor cells *in vitro* using a serial passaging protocol that mimics the end-replication problem. Aged progenitors demonstrated shorter telomeres (p<0.05), and reduced rates of cell proliferation, as marked by bromodeoxyuridine staining (p<0.001), with no changes in the ability of cells to differentiate into neurons or glia. RNA-sequencing and gene set enrichment analysis revealed an effect of cell ageing on gene networks related to neurogenesis, telomere maintenance, cell senescence and cytokine production. Downregulated transcripts showed a significant overlap with genes regulating cognitive function and risk for schizophrenia and bipolar disorder. Collectively, our results suggest that reductions in adult hippocampal neurogenesis, caused by telomere shortening, could represent a cellular mechanism contributing to age-related cognitive impairment and psychiatric disorder risk.

## Introduction

Patients with severe psychiatric disorders such as schizophrenia, bipolar disorder and major depressive disorder, have an average life expectancy approximately a decade lower than the rest of the population (1-3). This statistic primarily reflects the higher prevalence of comorbid age-related diseases, including coronary artery disease, diabetes and dementia, which contribute to early mortality (3, 4). Epidemiological findings such as these, have prompted researchers to consider whether faster ageing represents a core component to the pathophysiology of psychiatric disorders, or whether it represents the consequences of unhealthy lifestyles and stressful experiences, more common amongst those diagnosed with a psychiatric disorder (5, 6).

Telomere length is one biological marker which has been used to study rates of cell ageing amongst psychiatric disorder patients (7). Telomeres are DNA repeat structures found at the ends of chromosomes, which in humans, comprise of a six-nucleotide repeat sequence (TTAGGG) (8). Telomeres are vital for maintaining chromosomal stability (9) and have been shown to regulate a cell’s ability to replicate via mitosis (10). Telomere length gets shorter with each somatic-cell division due to the inability of DNA polymerase to fully replicate the 3’ end of the new DNA strand during DNA replication; a phenomenon known as the *end-replication problem* (11). Furthermore, telomerase activity (12), oxidative stress (13), genetic factors (14) and specific environmental factors, such as stress and exercise (15), have been found to moderate telomere length and the rate at which it shortens. When a telomere reaches a critically short length, the cell stops dividing and reaches a state of cellular senescence, often referred to as the ‘Hayflick limit’ (16). The Hayflick limit has been demonstrated in a variety of adult cell types *in vitro*, including fibroblasts (10), endothelial cells (17) and lymphocytes (18), whereby cells exhibit progressively shorter telomeres over increasing passages (or “population doublings”), a gradual reduction in their ability to proliferate, a reduced propensity to undergo programmed cell death, and a maladaptive proinflammatory phenotype (19). The reduced replicative capacity of ageing cells is hypothesized to contribute to age-related tissue-level pathology *in vivo*; as old, damaged cells, can no longer be replaced with new, healthy cells (20).

In the context of psychiatry, meta-analyses reveal that, in general, psychiatric disorder patients exhibit shorter leukocyte telomere lengths relative to unaffected individuals of equivalent ages, which could contribute to the increased burden of age-related disease (21-24). In addition to shorter telomere lengths, psychiatric disorder patients frequently exhibit neurological differences such as smaller hippocampi (25-27), and some studies have suggested a relationship between shortened telomere length and psychiatric disorder neuropathology (28-31).

The hippocampus is a brain structure important in cognition and mood regulation that is capable of adult neurogenesis due to the retainment of neural progenitors in the dentate gyrus; a specialised niche that allows progenitors to form new, functional neurons (32). Previous studies, including ours, have reported positive associations between peripheral telomere length and hippocampal volume (29, 33), and between telomere length and cognitive performance (29, 30), and it has been hypothesized that telomere length contributes to this association by moderating the rate of adult neurogenesis (34). This hypothesis is supported by various lines of evidence. First, chronological age is associated with both telomere length in hippocampal tissue, and rates of hippocampal neurogenesis (35-38). Consequently, telomere shortening in proliferating neural cell populations might be driving tissue-level differences in telomere length observed in the hippocampus. Second, reduced rates of hippocampal neurogenesis correlate with poorer performance in the Morris water maze task (39), and in visual pattern discrimination tasks (40), indicative of cognitive dysfunction; as well as reduced swim time in the forced-swim test, indicative of depression-like behaviour (41). Third, reductions in the expression of genes that regulate telomere length in the adult hippocampus in conditional knockout mouse models recapitulate age-related impairments to hippocampal neurogenesis, cognitive performance and behaviour, suggesting that telomere function is a key regulator of age-related changes to neurogenesis and cognition (42, 43). Despite these insights, to-date, there have been no studies investigating the impact of telomere shortening on *human hippocampal* progenitor cells, nor its downstream relationship to cognition and psychiatric disease. This is largely due to the inaccessibility of the dentate gyrus; the fact that there are no validated peripheral biomarkers or neuroimaging tools to assess rates of hippocampal neurogenesis *in vivo* in association with cognition and psychiatric conditions; and methodological challenges, like reliably immuno-staining post-mortem brain samples in large numbers (44).

The current study employed a systematic, multidisciplinary approach which aimed to model the effects of telomere shortening on human hippocampal neurogenesis, and subsequently its relationship to cognition and psychiatric disorder risk. First, we modelled telomere shortening using a serial passaging protocol that recapitulates the end-replication problem, in a human hippocampal progenitor cell line. We confirmed that reductions in telomere length were associated with lower levels of cell proliferation, without affecting the ability of cells to differentiate into neurons or glia. Second, complementary RNA-sequencing data revealed 3,281 transcripts which change in association with telomere shortening. Of these, the 1,594 downregulated genes strongly overlap with genes implicated in cognitive function and schizophrenia, and to a lesser extent bipolar disorder. Our work provides novel support for a multisystemic relationship, in which telomere shortening contributes not only to the heightened rates of age-related disease amongst psychiatric disorder patients, but its putative effects on hippocampal neurogenesis also implicate it as a mechanism important in the pathophysiology of psychiatric disorders themselves.

## Methods

### Human hippocampal progenitor cell line

An existing multipotent human fetal hippocampal progenitor cell line, HPCOA07/03 (ReNeuron, UK), was used to model human hippocampal neurogenesis *in vitro*, as used previously by our team (45-48). These cells proliferate in the presence of growth factors, and upon their removal, differentiate into neurons and glia. Our previous work demonstrates that after a seven-day differentiation protocol, cell populations primarily differentiate into doublecortin-positive immature neuroblasts, microtubule-associated protein 2 (MAP2)-positive neurons, and S100β-positive astrocytes (49). The remaining population maintains a neural progenitor cell phenotype. Immunocytochemistry has confirmed that MAP2-positive neurons co-stain for Prospero homeobox protein 1 (PROX1), which is a marker used to identify neurons from the dentate gyrus (46). For further details on the cell line and culture conditions for proliferating and differentiating cells, see Supplementary Information, S1-S2.

### Modelling telomere shortening via the ‘end replication problem’

As in other *in vitro* systems (17, 18, 50), we modelled telomere shortening by serially passaging cells. Serial passaging allows cell populations to double numerous times, and facilitates telomere shortening resulting from the end-replication problem (incomplete synthesis of chromosome ends during DNA replication).

Our experimental protocol utilised four cryovials of cells corresponding to four subcultures of cells (biological replicates), which were revived in separate T25 flasks and grown in proliferating medium. Once confluent, the cells were passaged onto a T75 flask under the same conditions, see Supplementary Information, S1 for further details. Cells growing at passage 21 were used to model *relatively* “young cells” of the adult hippocampus. Subsets of cells were then, either isolated and pelleted for DNA and RNA extraction (telomere length assessment and RNA-sequencing); seeded onto two 96-well plates for proliferation and differentiation assays (immunocytochemistry); or seeded onto a new T75 flask for subsequent serial passaging, see *Figure 1*. Passaging occurred once cells were 80-90% confluent (∼48 hrs), as confirmed by cell count. At each passage, cells were reseeded at a density of 2 x 10^6^ cells/T75 flask to ensure an approximately consistent number of population doublings across biological replicates. “Old cells” in our model corresponded to a subset of young cells that underwent eight subsequent passages, where again we isolated DNA and RNA, or submitted cells to a final proliferation or differentiation assay, *Figure 1*. A subset of cells was also collected at the midpoint of the assay (passage 25) to assess telomere length, which we refer to as “older cells”.

**Figure 1:**
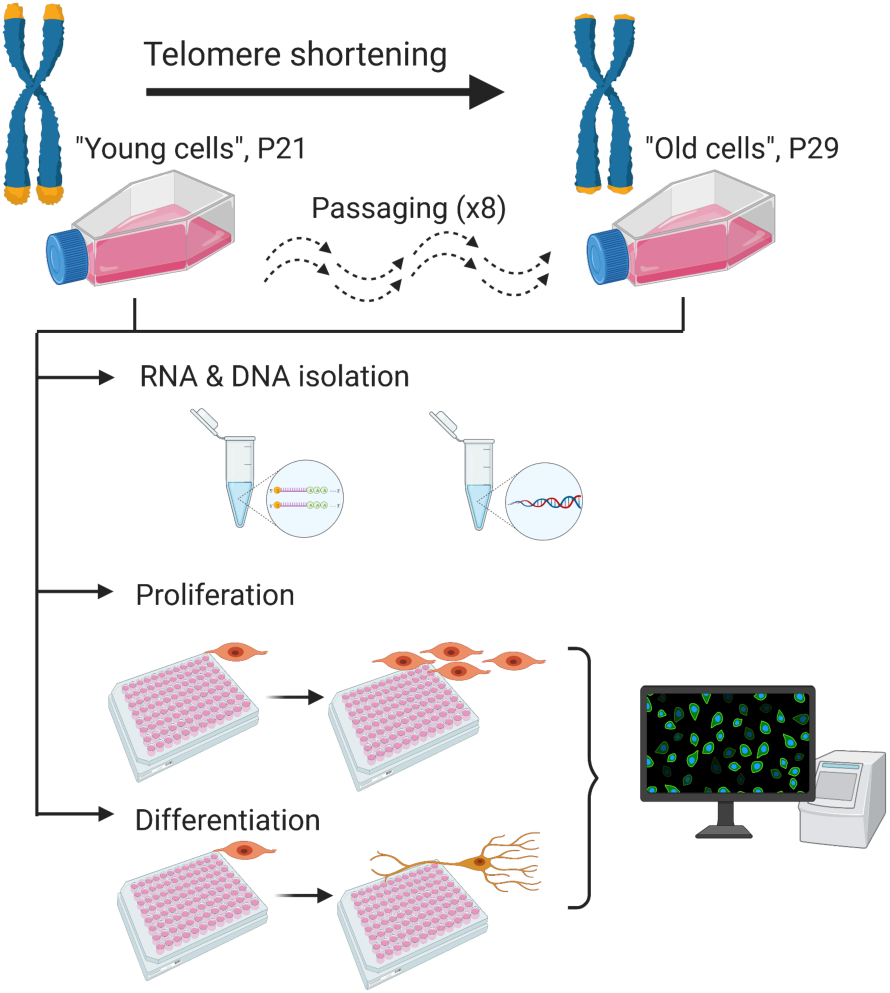
Protocol summary. A summary of our *in vitro* protocol, which considered mRNA levels, cell markers of proliferation, and cell markers of differentiation, in the context of telomere shortening.

### Proliferation and differentiation assays (Immunocytochemistry)

To compare differences in cell marker levels between young and old cells, we performed immunocytochemistry. Cells were seeded on laminin-coated 96-well plates at a density of 1.2 x 10^4^ cells/well in 100 μL of proliferating cell medium. Three technical replicates (wells) were generated in relation to each staining marker, alongside a negative staining control (no primary antibodies). Proliferating medium was replaced 24 hrs after seeding and replaced with fresh medium. Cells were then grown for 48 hrs after which they were fixed using a 4% paraformaldehyde solution. For three of the wells, we included a Bromodeoxyuridine (BrdU) incorporation step, consisting of a 10 μM BrdU treatment for the last 4 hrs prior to fixing. Subsequently, immunocytochemistry was used to assay proliferation markers including BrdU and Ki67, and the apoptosis marker caspase-3 (CC3).

For cells submitted to the differentiation protocol, they underwent the same conditions as the proliferating cells, but after 48 hrs, instead of fixation, the cells were washed twice with differentiation medium in order to remove remaining growth factors and cell debris. Cells were then incubated for a further seven days with no media changes, as previously described (51). After seven days, cells were fixed in order to assay cellular markers. Cell markers assessed included the neuronal markers, doublecortin and microtubule-associated protein 2 (MAP2), the astrocyte marker, S100β, and apoptotic marker, CC3. Quantification of immunostaining was performed using the unbiased and semi-automated high-throughput Thermo Scientific Cell-Insight CX5 High Content Screening Platform (Thermo Scientific, Massachusetts, USA). See Supplementary Information, S3-S4 for further details, and Table S1 for a list of primary and secondary antibodies.

### Nucleic acid extraction

Approximately 50% of the cell suspension obtained during passaging was pelleted and stored at −80°C at the beginning (passage 21; young cells), midpoint (passage P25; older cells) and end of experiments (passage P29; old cells). DNA and RNA were extracted using the Qiagen AllPrep DNA/RNA/Protein Mini Kit (Qiagen, Hilden, Germany; #80004). The standard manufacturer’s protocol was followed to extract DNA and RNA from each cell pellet, with the inclusion of an additional ethanol precipitation at the end, for nucleic acid purification. All DNA samples had 260/280 ratios of between 1.7 and 1.9, and RNA samples had 260/280 ratios of 1.95 to 2.1, tested using the Nanodrop D1000 (Thermo Scientific, Wilmington, DE), indicating samples were of good purity. All RNA samples had integrity numbers (RINs) greater than 9, as tested using the Agilent 2100 Bioanalyzer (Agilent Technologies, Berkshire, UK).

### Telomere length measurements

Relative telomere length was quantified using DNA samples and a modified version of the quantitative Polymerase Chain Reaction (qPCR) protocol described by Cawthon and colleagues (52), as used by our lab previously (7, 14, 29, 53). First, the protocol assayed the telomere variable repeat region (TTAGGG), and the cycle threshold (C*t*) required to reach a predetermined level of fluorescence: this correlated with the number of telomere repeats present in the individual samples. Second, a single-copy gene (albumin) was assayed in parallel, except the C*t* now correlated with the number of copies of the genome in that individual DNA sample. Finally, a telomere-to-single-copy-gene ratio was used to determine relative telomere length, where the number of telomere repeats in each sample was corrected for the total number of copies of the genome in the individual DNA sample being tested; see Supplementary Information, S5.

### RNA-sequencing

Library preparation and RNA-sequencing was performed at The Genomic Centre, King’s College London. Briefly, total RNA samples were submitted to a DNAse treatment using the DNA-free™ DNA Removal Kit (Invitrogen, California, USA). Subsequently, 300 ng of total RNA from each sample was submitted for ribosomal RNA depletion using the NEBNext rRNA Depletion kit (New England Biolabs, Massachusetts, USA), and RNA-seq libraries were constructed using the NEBNext Ultra II Directional RNA Library Prep Kit for Illumina. The samples were sequenced in a HiSeq 4000 sequencing system (Illumina).

Raw reads were downloaded and processed using Trimmomatic 0.38 (54), to prune low quality bases (leading/trailing sequences with phred score < 3, or those with average score < 15 every four bases), or reads below 36 bases in length. Trimmed reads were pseudoaligned to the human reference genome GRCh38 using kallisto (55). The package tximport (56) was used to import kallisto output files into DESeq2 (57), summarizing transcript-level information to gene-level expression counts utilizing gene information from the Ensembl Release 99, imported using biomaRt (58). Differential expression analysis was performed in young versus old cells (N = 4 biological replicates per condition) using the Wald test in DESeq2, controlling for biological replicates. Log2 fold-changes were shrunk using apeglm (59), and the false discovery rate (FDR) correction was used to control for multiple comparisons. Gene expression differences were considered significant if P_FDR_ < 0.05.

### Gene Ontology Enrichment

To understand which biological mechanisms were being upregulated and downregulated within our cell model in association with telomere shortening, we separately entered genes (P_FDR_ < 0.05) showing an increase in expression, and those showing a decrease in expression, into FUMA (60). The GENE2FUNC tool performs a gene set enrichment analysis, where gene sets were defined by The Molecular Signatures Database (MSigDB). We included all transcripts surviving DESeq2’s internal filtering criteria as our background list (i.e. all genes expressed in the samples). Multiple testing correction was performed using the FDR method, and GO terms were considered significant if P_FDR_ < 0.05.

### Gene set enrichment analysis

We tested for a genetic overlap between genes affected in our cell ageing model and those implicated in major depressive disorder (61), bipolar disorder (62), schizophrenia (63) and general cognitive function (64), using publicly available GWAS summary statistics. As previously (46), we separately tested genes which were downregulated and upregulated in our cell model. Gene set enrichment was performed in MAGMA (65), using a 10 kb annotation window around genes, and after removing the region encompassing the major histocompatibility complex (chromosome 6, 25-34 Mb). MAGMA calculates gene-level enrichment by generating a gene-wide statistic from GWAS summary statistics, adjusting for gene size, variant density, and linkage disequilibrium using the 1000 Genomes Phase 3 European reference panel. Subsequently, MAGMA was used to perform a competitive test of gene set association. This competitive test of association calculates the enrichment of how over-represented the genes in our gene sets are, in relation to those implicated in each trait, and relative to other gene sets of similar size across the genome. We corrected the p-values output by MAGMA by the number of traits (four) and the number of gene sets (two) tested, using the Bonferroni method.

### Statistical analyses

When assessing telomere length changes in association with passaging, ANOVA were used, followed by Tukey’s multiple comparison test. Cell marker differences between young and old cells were determined using two-tailed t-tests, followed by a Bonferroni correction to account for the number of markers assayed per condition. RNA-sequencing data and downstream analyses were performed as described above.

### Figure generation

Figures were created using Prism7 (GraphPad, San Diego USA), the EnhancedVolcano package in R (66) and BioRender (Toronto, Canada).

## Results

### (i) Hippocampal progenitor cells demonstrate sustained telomere loss in response to passaging and the end-replication problem

We compared relative telomere length from young cells (P21), older cells that correspond to the midpoint (P25) of the experiment, and old cells (P29) from the end of our cell protocol. ANOVA revealed significant differences in telomere length between the groups (F(4, 9) = 9.349, P = 0.013), whereby telomeres were significantly shorter in older cells (Tukey’s post-hoc test, p < 0.05), and old cells (p < 0.05) relative to young cells, Figure 2. This result indicates a sustained loss of telomere length in response to passaging.

**Figure 2:**
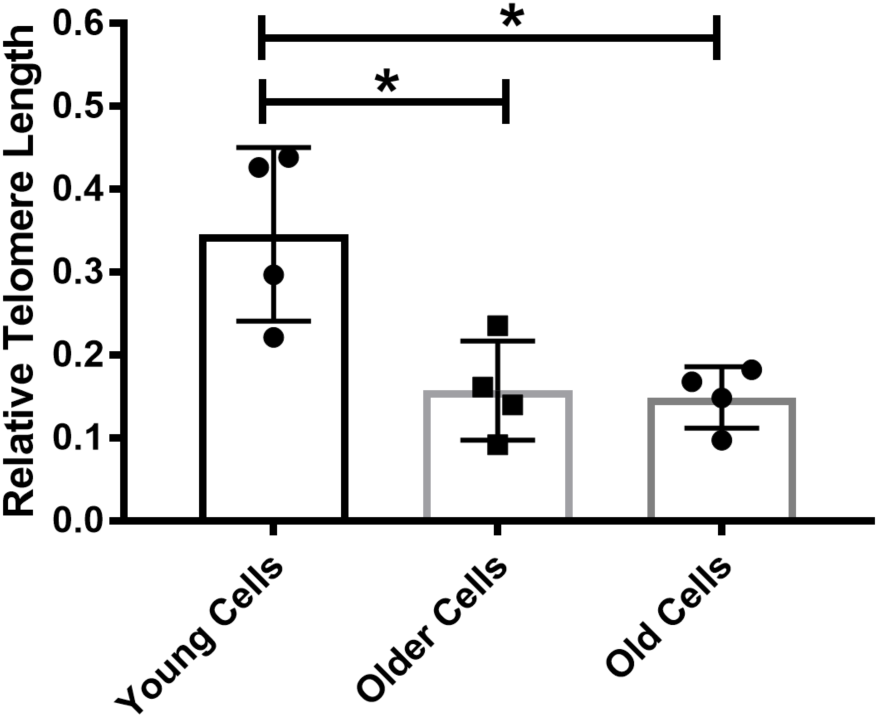
Telomeres are shorter in old hippocampal progenitor cells. This bar chart shows the relative telomere length in young cells (P21), older cells (P25) and old cells (P29). There is a significant reduction in telomere length in both the older and old cells, relative to young cells. Group differences were detected using a one-way ANOVA with Tukey’s post-hoc test. Significant differences were considered when P < 0.05, indicated by *. N = 4 for all groups.

To complement this finding, we confirmed that telomere shortening was likely being driven by the end-replication problem, as opposed to changes to the telomerase enzyme (another critical telomere regulator) during passaging. *hTERT* codes for the catalytic subunit of the telomerase enzyme and is tightly controlled, and closely associated with enzyme activity (67). We performed a quantitative PCR to assess differences in the expression of telomerase reverse transcriptase (*hTERT*) in old and young cell, and found consistently low levels of *hTERT* expression, which did not differ between groups (P > 0.05); see Supplementary Information, S6.

### (ii) Old hippocampal progenitor cells exhibit decreased cell proliferation

Cells demonstrated a significant reduction in proliferation in association with telomere shortening, Figure 3. Old cells showed significantly lower levels of proliferation relative to young cells, as marked using BrdU staining (two-sample t-test, t(6) = 6.663, p < 0.001), a difference which remained significant after correcting for the number of cell markers tested (p < 0.05). This difference was also supported by quantification of the proliferation marker Ki67 (two-sample t-test, t(6) = 2.959, p = 0.025). We also observed a lower percentage of old cells stained with CC3 (indicative of lower rates of cell death), relative to young cells (two-sample t-test, t(6) = 2.481, p = 0.047), however this effect did not survive multiple testing correction (p > 0.05); representative CC3 staining is shown in Figure 4.

**Figure 3:**
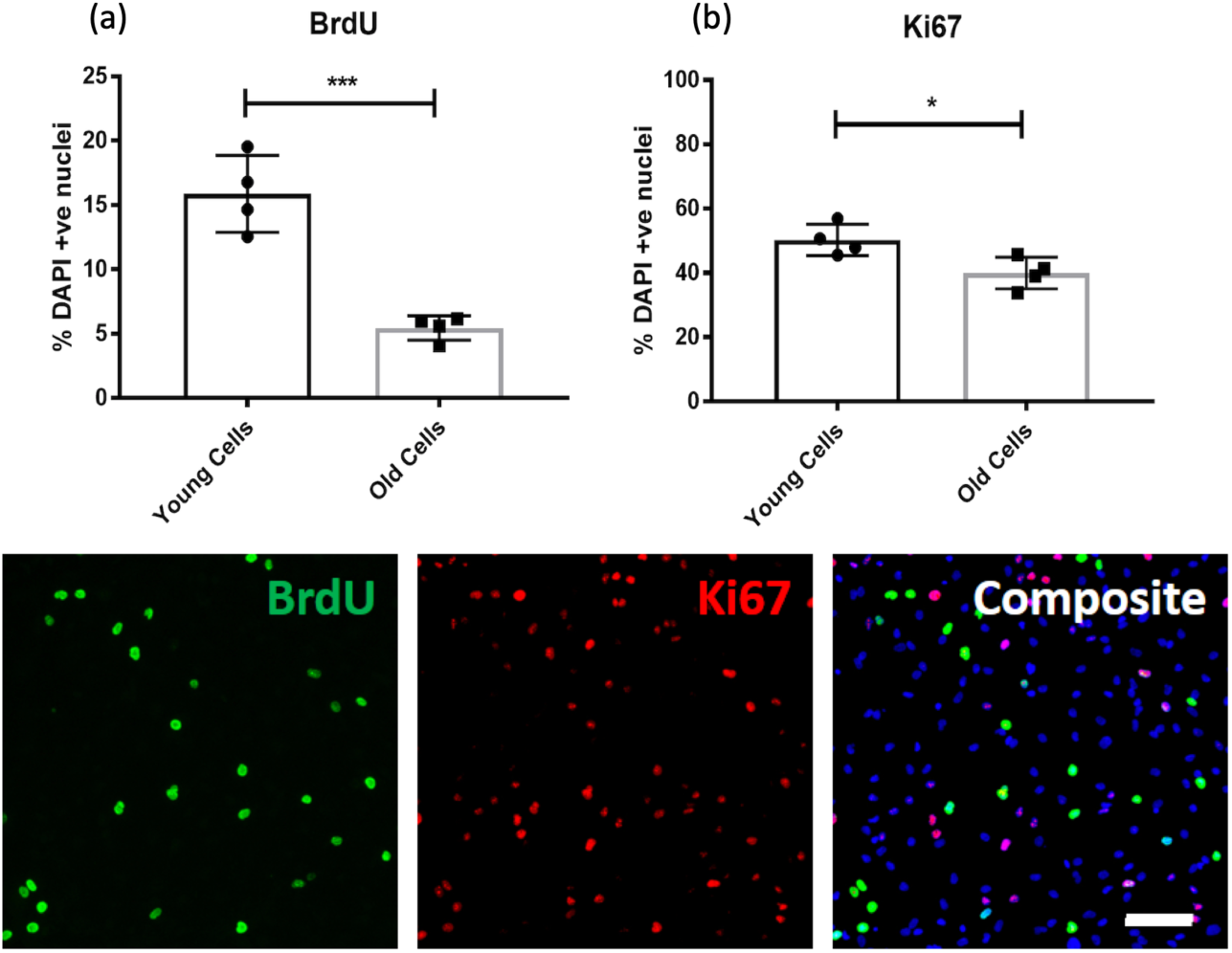
Proliferation rates are lower in old hippocampal progenitor cells. Bar charts (top) show the percentage of BrdU (a) and Ki67 (b) positive cells relative to the percentage of DAPI stained nuclei (y-axis) in young cells and old cells (x-axis). Each data point represents one biological replicate (N = 4). *** represents an uncorrected p < 0.001, and * represents an uncorrected p < 0.05. Each of the images (below) are representative of a field of immuno-stained cells, taken using a 10X objective with the CellInsight High Content Screening Platform. Each composite image includes the nuclear marker DAPI in blue. Scale bar = 100 μm.

**Figure 4:**
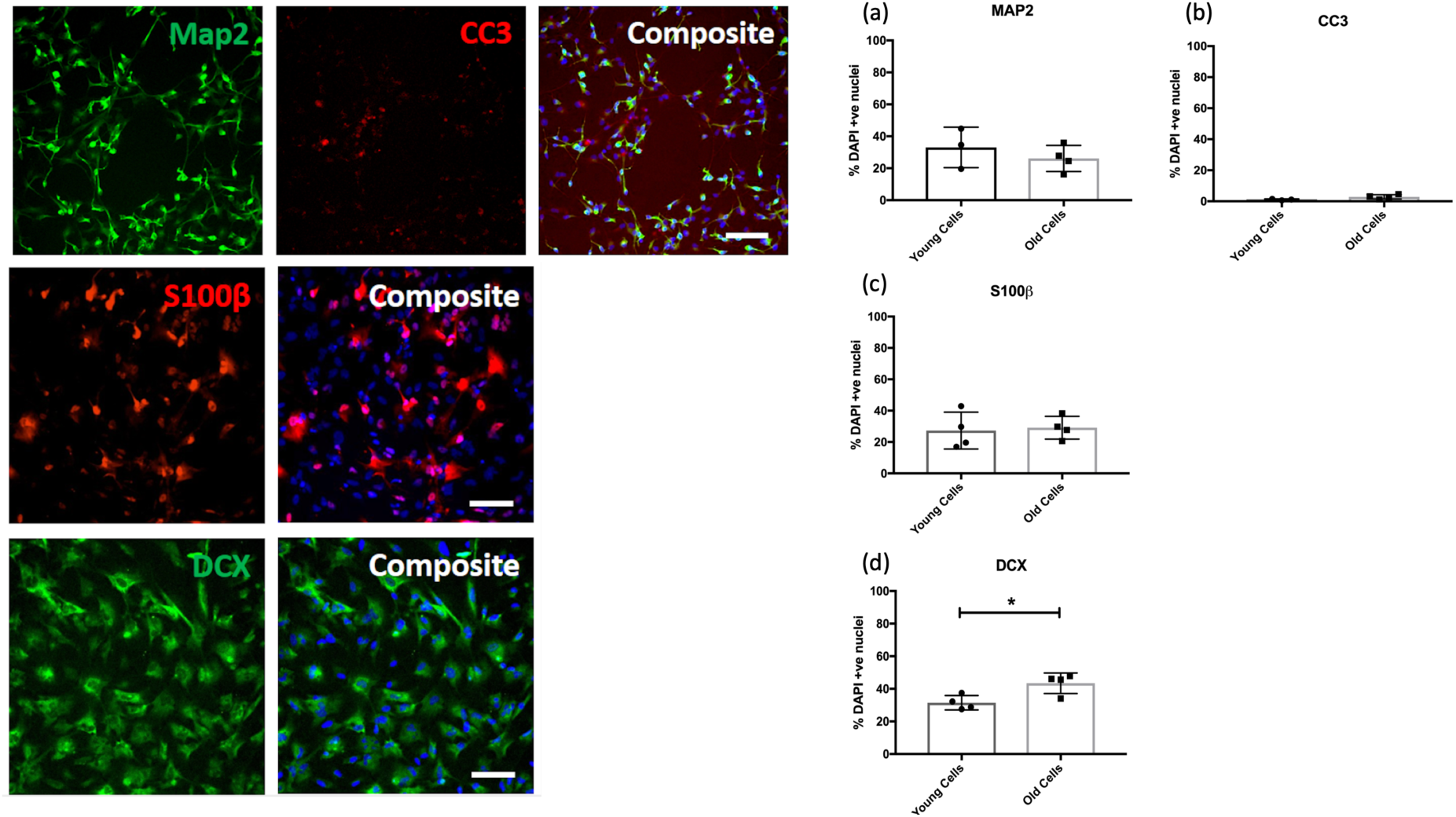
There are no differences in the rates of cell differentiation in old cells. The bar charts (right) show the percentage of MAP-2 (a), CC3 (b), S100β (c) and doublecortin (DCX) (d) positive cells relative to the percentage of DAPI stained nuclei in young cells and old cells. Each data point represents one biological replicate (N = 4). * represents an uncorrected p < 0.05. Each of the images (left) are representative of a field of immuno-stained cells, taken using a 10X objective with the CellInsight High Content Screening Platform. Each composite image includes the nuclear marker DAPI in blue. Scale bar = 100 μm.

### (iii) Old hippocampal progenitor cells do not show differences in their rate of cell differentiation

We observed no differences (p > 0.05) in markers pertaining to glial cells (S100β), cell death (CC3), or more mature neurons (MAP-2), between young and old cells. There was a small increase in the number of doublecortin positive neurons observed in old cells relative to young cells (two-sample t(6) = 3.097, p = 0.021), though this difference did not survive multiple testing correction (p > 0.05), Figure 4.

### (iv) Cell ageing is associated with vast transcriptional changes related to neurogenic processes, cellular senescence and inflammation

We identified 3,281 transcripts which were differentially expressed in old cells relative to young cells (P_FDR_ < 0.05), Figure 5a. We found that 1,594 genes were downregulated in old cells. These genes were broadly associated with neurogenesis and nervous system development. Furthermore, enrichment amongst canonical pathways revealed an over-representation of genes related to telomeres and cell senescence, Figure 5b.

**Figure 5:**
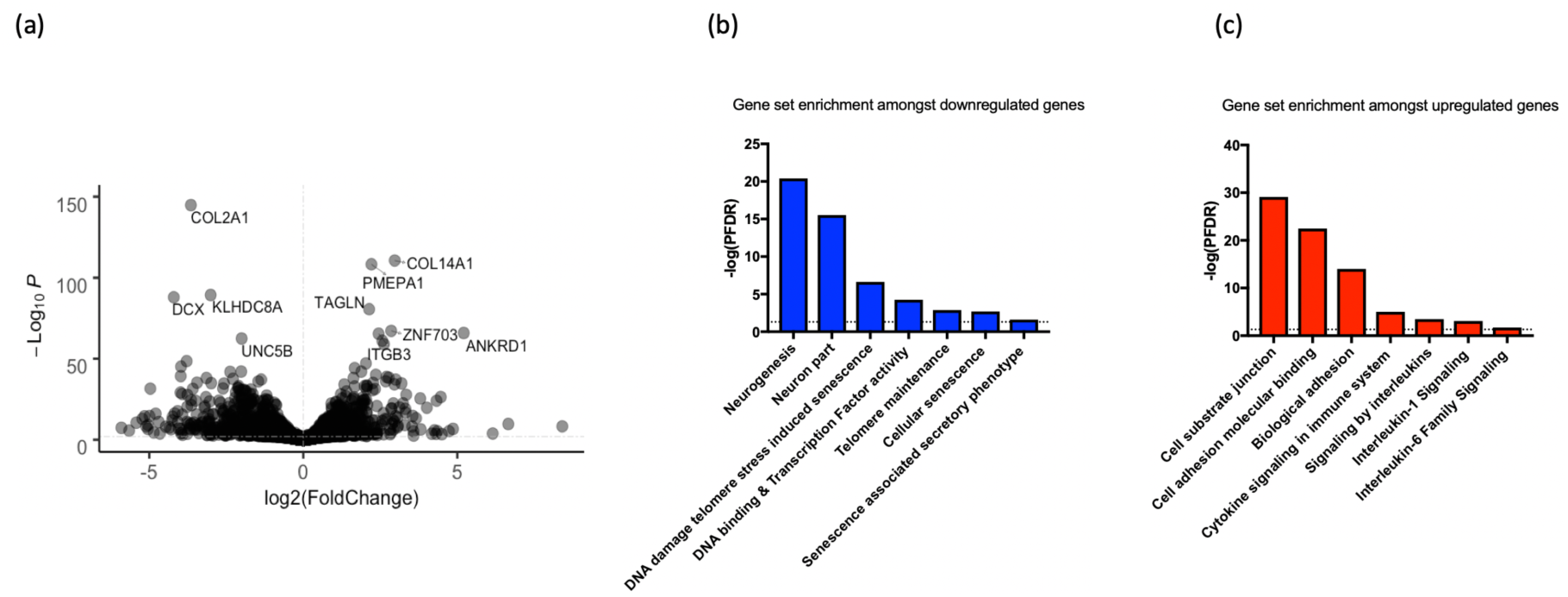
Differentially expressed genes in old cells relative to young cells, and examples of gene sets implicated in cell ageing. (a) A volcano plot summarizing the RNA-sequencing results, where log2(Fold change) is shown on the x-axis, and the strength of the association given by −Log_10_(P), is shown on the y-axis. (b) Examples of gene sets which significantly overlap with the downregulated genes in our cell model (c) Examples of gene sets which significantly overlap with the upregulated genes in our cell model. All gene sets represent those which surpassed a false discovery rate correction, P_FDR_ < 0.05, in our enrichment analysis, as marked by the dashed line.

**Figure 6:**
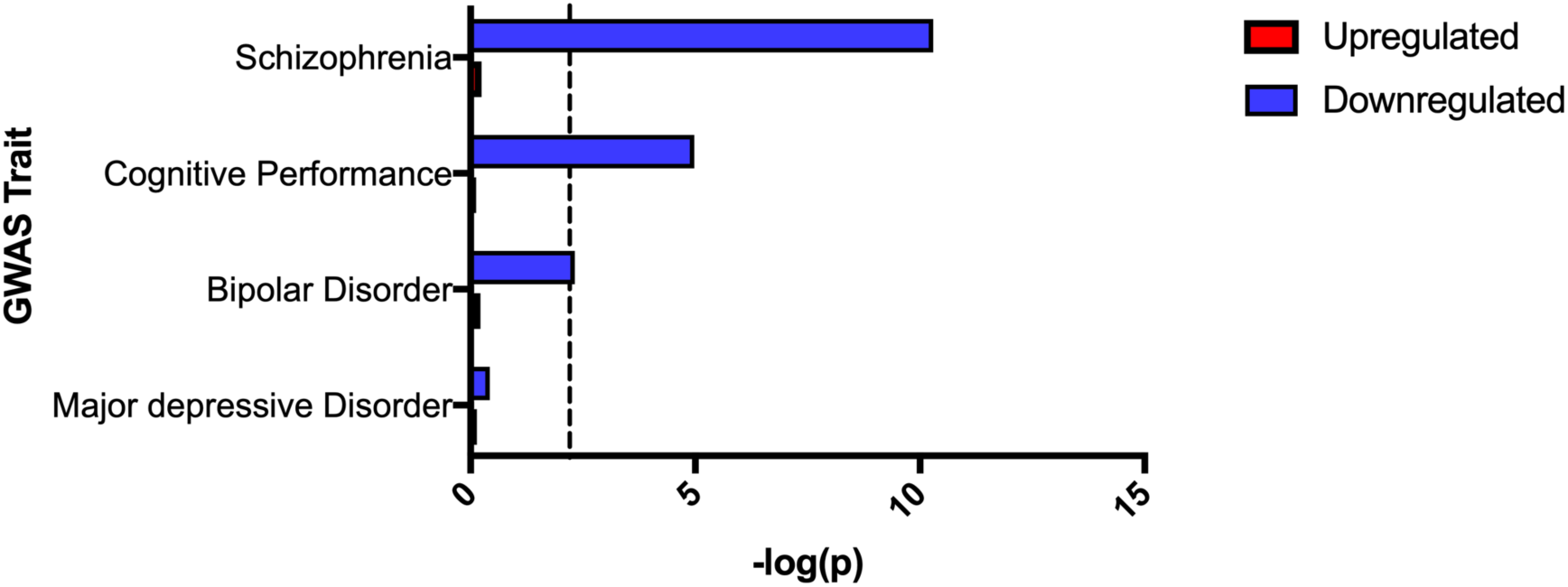
Downregulated genes in response to cell ageing are associated with cognitive performance and psychiatric disorder risk. A bar plot showing the genetic overlap between various traits as assayed by GWAS (y-axis) and genes either upregulated or downregulated in association with cell ageing. The strength of the association is shown on the y-axis (-log(p)). The dashed line represents the threshold of significance (corrected for the number of tests).

1,687 genes were upregulated in old cells, and these genes were associated with cell adhesion. Enrichment amongst canonical pathways revealed an over-representation of genes related to inflammation and cytokine signalling, Figure 5c. In agreement with previous work relating to the senescence associated secretory phenotype, amongst the upregulated transcripts we observed increased expression of the cytokine regulator, Nuclear Factor Kappa B (*NFKB1*) and increased interleukin-6 (*IL6*) levels (PFDR < 0.05) (19). See Supplementary Datasets for full results.

### (iv) Genes downregulated in old cells overlap with those implicated in cognitive performance, schizophrenia, and bipolar disorder risk

We found a significant overlap between genes which were downregulated in old cells (P_FDR_ < 0.05) and those implicated in: schizophrenia (β = 0.207, SE = 0.032, P = 5.105 x 10^−11^); bipolar disorder (β = 0.071, SE = 0.027, P = 0.005); and cognitive performance (β = 0.138, SE = 0.037, P = 1.057 x 10^−5^), based on GWAS results.

To ensure this result was not being inflated by the relatively large size of our gene set (n=1,594), or by an overrepresentation of genes related to tissue type, we performed a sensitivity analysis. From the transcripts surpassing our multiple testing threshold, we selected 500 which exhibited the highest fold change. Additionally, when mapping SNPs to genes, we removed any genes which were not expressed in our cell line from the genomic annotation. When repeating gene-set enrichment analysis, we confirmed that downregulated transcripts remained significantly associated with each trait (P ≤ 0.01). We found no significant overlap between downregulated genes and those implicated in major depressive disorder, nor between upregulated genes and any of our traits.

Previous evidence suggests that telomere shortening and reduced levels of hippocampal neurogenesis are specific to the adult brain (35-37), but schizophrenia risk and cognitive function are also known to be affected by prenatal neurodevelopmental factors (68). We performed a complementary experiment using fetal brain samples in order to clarify whether telomere shortening occurs in the human brain during early development, and therefore, whether or not our cell model could also be recapitulating a neurodevelopmental cell process. We assessed telomere length in fetal brain samples collected from various post-conception days (75-161 days), which is a measure that positively correlates with the number of cell divisions (69). We did not find a significant correlation between telomere length and post-conception days (p > 0.05), Supplementary Information, S7. This suggests that the effect we observe in our study, would be unlikely to occur in early neurodevelopment, further supporting the specificity of this process to the postnatal brain.

## Discussion

Our study used serial passaging, an established method for modelling telomere shortening via the end replication problem, in human hippocampal progenitor cells. Aged cells demonstrated reduced rates of cell proliferation, as determined by BrdU and Ki67 immunostaining. Telomere shortening was further accompanied by changes to 3,281 transcripts, whereby downregulated genes overlapped with those implicated in cognition, schizophrenia, and bipolar disorder.

The findings presented here support previous research performed in other cell model systems that reveal an intricate relationship between telomere length and cell replicative capacity related to the end-replication problem (17, 18, 50). Our gene expression data further demonstrates an enrichment of transcript changes related to cell senescence, the senescence associated secretory phenotype, and interleukin-6 signalling. This is in agreement with previous research (19), and validates that our model is capturing meaningful biological changes related to telomere shortening and cell ageing. It also suggests that the decreased proliferation and nominally decreased cell death we found in old progenitors, likely relates to early signs of cell senescence, i.e. a reduced number of new, healthy cells, and an accumulation of old, unhealthy cells that instead of apoptosing, demonstrate a proinflammatory phenotype.

In the context of neurogenesis, our findings extend prior work in animals and post-mortem brain which have shown a significant decline in markers of cell proliferation in the hippocampus in association with age (70, 71), by further implicating telomere shortening as one potentially important age-related cellular mechanism. In contrast to some work (36, 72), we did not observe a robust effect of telomere shortening on rates of neuronal differentiation, as marked by doublecortin and MAP-2 in our cell staining data. However, amongst proliferating cells there was a strong downregulation of doublecortin at the transcript level in association with telomere shortening (*Fig. 5a*), which might suggest that there are very early reductions in the rates of differentiation, which were not observable after the seven-day differentiation protocol used in our model.

Genetic and association studies have inferred a causal relationship between telomere length and cognition (73), but no study to-date, has demonstrated a cellular mechanism. Our work supports these findings and demonstrates that telomere shortening affects genes which regulate general cognitive function, in association with reduced hippocampal progenitor cell proliferation. Our cell model also implicates a relationship between telomere shortening in human hippocampal progenitors and schizophrenia and bipolar disorder risk. Both schizophrenia and bipolar disorder share a common aetiology (74), and patients frequently exhibit smaller hippocampal volumes (25-27) and general cognitive deficits relating to attention, working memory, verbal learning and memory and executive functions (75). In the context of our results, it is possible that telomere-induced reductions in hippocampal neurogenesis represent one mechanism that contributes to cognitive dysfunction amongst patients.

Major depressive disorder patients also exhibit cognitive dysfunction, though to a lesser and more specialised extent than schizophrenia patients (76), however, we did not find an enrichment of major depressive disorder genes in association with telomere shortening in our cell model. This could relate to a difference in the extent and nature of the cognitive dysfunction, or the less heritable nature of depression and more penetrant influences of environmental stressors in its aetiology (77). The latter possibility is supported by a recent study in the UK Biobank which suggests that early life stress can amplify the penetrance of genetic risk signals in predisposing to major depressive disorder (78). In addition, it is supported by animal and *in vitro* studies which suggest that the downstream effects of stress, particularly increased cortisol levels and glucocorticoid receptor activation, might moderate depression risk by affecting the proliferation of hippocampal progenitors (41, 49). Consequently, it is possible that the genes regulating responses of hippocampal progenitor cells to cortisol exposure, for example, could be more pertinent to the aetiology of major depressive disorder (79).

Our results also suggest that environmental factors with the potential to prevent premature telomere shortening may have positive influences on hippocampal neurogenesis and cognitive function. Intriguingly, environmental interventions shown to reduce the rate of cognitive ageing (80), including exercise and energy restriction, have positive effects on both peripheral telomere length in humans (15, 81, 82) and hippocampal neurogenesis in animal models (83, 84). Furthermore, diets rich in omega-3 fatty acids and antioxidants, have been associated with both increased rates of hippocampal neurogenesis and longer telomeres (32, 85, 86), as have some drugs, such as resveratrol and lithium (14, 87-89). This implies that environmental, dietary and pharmacological interventions could be useful in systematically targeting premature cell ageing and reduced hippocampal neurogenesis, and in doing so help to maintain mental (and physical) health, of individuals as they age. In the case of schizophrenia, studies have already shown the beneficial influences of exercise on health and cognition (90), suggesting these interventions more broadly, may have benefits for patient groups that demonstrate faster rates of cell ageing.

Despite the promising evidence shown here, there are a number of limitations to our study which should be acknowledged. First, we are measuring changes to telomere length and its effect on hippocampal neurogenesis using a cell model system acquired from a single donor. It is possible that individuals from different genetic backgrounds may respond differently to the effects of telomere shortening (14), and as such, future work utilising induced pluripotent stem cell models from a range of donors would be beneficial in validating and extending our work. Second, although our cell model demonstrated sustained telomere shortening and signs of cellular senescence, the effect was modest, with some old cells continuing to proliferate (10). It is likely that a longer culture protocol will provide greater insights into the progressive and longer-term consequences of telomere shortening in hippocampal cells, and this is an important future consideration. Third, the cell model attempts to understand the specific effects of telomere shortening on adult hippocampal neurogenesis, in order to better understand age-related changes to the brain. However, alongside changes in telomere length, there may also be independent changes to the epigenetic landscape (91), so future work will also be needed to further differentiate the causative effects of telomere shortening (e.g. via *TERT* knockout), from other age-related cell changes. Furthermore, our model makes use of fetal-derived neural progenitors, which although studies suggest do recapitulate the functions of adult neural stem cells, they may not do so entirely (92). Moreover, our work in fetal brain indicates that despite the importance of early neurodevelopmental factors in the etiology of schizophrenia (68), telomere shortening does not appear to occur during early brain development. Consequently, further work will be needed to confirm at what stages in postnatal life telomere shortening impacts on hippocampal neurogenesis and becomes important in the pathophysiology of psychiatric disease.

This work extends our understanding of the age-related molecular mechanisms influencing hippocampal neurogenesis, and implicates telomere shortening as an important process associated with psychiatric disorders and cognitive function. It also provides novel support for a multisystemic relationship, in which telomere shortening contributes not only to the heightened rates of age-related disease amongst psychiatric disorder patients, but its putative effects on hippocampal neurogenesis also implicate it as a mechanism important in the pathophysiology of psychiatric disorders themselves. Future work should consider whether leukocyte telomere length could act as a useful peripheral biomarker to estimate changes in the rates of hippocampal neurogenesis *in vivo*, and whether environmental interventions could be used to simultaneously prevent premature telomere shortening and promote hippocampal progenitor proliferation, as we age.

## Acknowledgments

This work was funded by a Medical Research Council Skills Development Fellowship (MR/N014863/1) and a Psychiatry Research Trust Grant (grant reference: 92 Branthwaite) awarded to T.R.P., as well as by Medical Research Council funding (MR/N030087/1) awarded to S.T. A.B.P. is funded by a Rayne Foundation PhD studentship and by the National Institute for Health Research (NIHR) Mental Health Biomedical Research Centre, South London and Maudsley NHS Foundation Trust and King’s College London. The views expressed are those of the authors and not necessarily those of the NHS, the NIHR or the Department of Health and Social Care. The funding sources had no role in the study the design, in the collection, analysis and interpretation of data, in the writing of the report and in the decision to submit the article for publication. The fetal material was provided by the Human Developmental Biology Resource (http://www.hdbr.org). Ethical approval for the HDBR was granted by the Newcastle & North Tyneside 1 Research Ethics Committee under the reference 18/NE/0290. We would like to thank Dr Nick Bray for his constructive feedback on this work and for providing us with access to the human DNA samples used here.

## Author contributions

A.B.P., S.T. and T.R.P. designed research; A.B.P., D.M.S., E.C.H., R.R.R.D, S.T. and T.R.P. carried out or led laboratory work; D.F.N. contributed knowledge and critical input; A.B.P., R.R.R.D, and T.R.P. analysed the data; A.B.P. and T.R.P. wrote the paper with the assistance of all authors.

