## Supplementary Information for "The impact of telomere shortening on human hippocampal neurogenesis: Implications for cognitive function and psychiatric disorder risk"

**S1: Hippocampal Progenitor Cell Line:** We utilised a multipotent human fetal hippocampal progenitor cell line (HPCOA07/03; ReNeuron, UK), as in our previous work (1-4). The cell line was derived from first-trimester, female, foetal hippocampal tissue following medical termination, and in accordance with UK and USA ethical and legal guidelines, and obtained from Advanced Bioscience Resources (Alameda CA, USA).

HPC0A07/03C cells were *conditionally* immortalised by introducing the c-myc-ERTAM transgene, whereby upon activation of the modified estrogen receptor with 4-hydroxy-tamoxifen (4-OHT), the oncogene c-myc becomes activated and maintains high levels of proliferation in culture, with the aid of the growth factors, epidermal growth factor (EGF) and basic fibroblast growth factor (bFGF) (5). Upon removal, cells differentiate into doublecortin (DCX)-positive immature neuroblasts, microtubule-associated protein 2 (MAP2)-positive neurons, and S100β-positive astrocytes. The remaining population maintain a neural progenitor cell phenotype. Immunocytochemistry has confirmed that MAP2-positive neurons co-stain for Prospero homeobox protein 1 (PROX1), which is a marker used to identify neurons from the dentate gyrus (2).

Due to the established effect of c-myc activation on maintaining high telomerase activity and telomere length (6), for the purposes of these experiments, we eliminated 4-OHT from our culture at the point of cell reactivation (passage 16), and maintained cells in a proliferating state using EGF and bFGF only. To ensure thorough removal of 4-OHT and to eliminate any subtle downstream effects of 4-OHT removal on cells, we grew them for five passages without 4-OHT, prior to the initiation of experiments. Across these five passages we did not observe any significant changes in the length of time required for cells to reach confluency, indicating the absence of a global, incremental impact of 4-OHT removal on cell number. Furthermore, consideration of *hTERT* expression (a telomerase gene which strongly correlates with its activity (7)) in old and young cells indicated consistently low levels of telomerase activity in cells, suggesting residual c-myc activation was unlikely to be influencing telomere length, see S6.

**S2: Cell Culture conditions:** Cells were cultured in monolayer on Nunclon flasks (Thermo Fisher Scientific, Massachusetts, USA; #156367/156499), coated with 23 μg/ml laminin from Engelbreth–Holm–Swarm murine sarcoma basement membrane (Sigma, Missouri, USA) in phosphate buffered saline (Gibco, Massachusetts, USA; #L2020) for 1–24 hrs to aid cell adhesion. When passaging, media was aspirated and cells were exposed to warm accutase (Sigma, #A1110501) to lift them from monolayer, they were then resuspended in warm medium, and washed twice via centrifugation at 900 rpm for 5 minutes with subsequent resuspension. Cells were maintained in chemically-defined proliferation medium and incubated at 37 °C, 5% $CO_2$ and sufficient humidity. Proliferation medium was composed of Dulbecco's modified Eagle's Medium Nutrient Mixture F-12 Ham (DMEM-F12; Sigma–Aldrich, #D6421 or #D6434) supplemented with 0.03% human albumin solution (Zenalb, #20), 100 μg/ml human apo-transferrin (Sigma, #T1147), 16.2 μg/mL human putrescine DiHCl (Sigma, #P5780), 5 μg/mL human recombinant insulin (Sigma, #I9278), 60 ng/mL progesterone (Sigma, #P8783), 2 mM l-glutamine (Sigma, #G7513) and 40 ng/mL sodium selenite (Sigma, #S9133). 10 ng/mL human bFGF (Peprotech, New Jersey, USA; #AF 100-15-500) and 20 ng/mL human EGF (Peprotech, #EC 100-18B) were including in proliferating cell media only.

**S3: Cell fixing & immunocytochemistry**

At the end of the proliferation/differentiation protocols, the culture medium was aspirated and cells were washed with warm cell media to remove any cell debris. The cells were then fixed with 50 µl/well of freshly thawed 4% paraformaldehyde (PFA, Alfa Aesar 43368) in phosphate buffered saline (PBS) and incubated for 20 mins at room temperature and in the dark. The PFA was removed after 20 mins and the fixed cells were washed three times with PBS. The plates were then stored in 0.05% Sodium Azide (Sigma, S8032) in PBS, wrapped in parafilm (Sigma, P7793) at $4^{O}$C.

Immunocytochemistry (ICC) was carried out in order to analyse the fate, viability and health of these cells, based on morphological and protein localisation quantification. For BrdU cells only, following fixation, the cells were incubated with 2N hydrochloric acid for 40 mins in order to denature the DNA strands. Cells were then neutralised with 0.1 M sodium borate buffer for 10 mins and washed with PBS.

For all cells, blocking solution comprised of 5% normal donkey serum (Sigma, D9963) and 0.3% Triton X-100 (Sigma, T9284) in PBS was used to block cells for 1 hr at room temperature. Primary antibodies were then diluted to the appropriate concentration in blocking solution and added to the cells for overnight incubation at $4^{O}$C. Antibodies were carefully chosen to identify cells at varying degrees of proliferation, differentiation and apoptosis, see *Table 1*. First, the cells were washed with PBS and incubated with blocking solution for 30 mins. During these 30 mins, secondary antibodies were diluted in blocking solution at the appropriate concentrations and added to cells for 2 hrs. Cells were then washed twice with PBS and the nuclei were stained with 300 nM DAPI for 5 mins. Following this, cells were washed twice with PBS and stored in 0.05 % sodium azide in PBS at $4^{O}$C, wrapped in tin foil to block sunlight and prevent photobleaching.

Every plate was designed to contain unstained control cells which were used as a benchmark for background fluorescence in downstream analyses. Antibody combinations were kept the same for all experiments (Table 1) and representative images are shown in the main text.

**S4: Quantification of cell morphology and protein localisation**

Quantification of cell antibody staining was performed using an unbiased and semi-automated high-throughput Thermo Scientific Cell-Insight CX5 High Content Screening Platform (Thermo Scientific, Massachusetts, USA) alongside the native HCS Studio Cell Analysis Software (Thermo Scientific). Separate protocols were developed for each antibody using the Cell Health Profiling BioApplication, which first identifies cells based on their nuclear staining and then quantifies the intensity of fluorescent staining in user-defined regions. Once a protocol was developed, it was kept constant across all the other plates and biological replicates. On rare occasions the parameters were changed to account for differences in the brightness of immunofluorescence between biological replicates.

| *Primary Antibody* | *Dilution* | *Supplier* | *Catalogue #* | *Marker usage* |
|---|---|---|---|---|
| Rat anti-**BrdU** | 1:500 | Serotec | OBT0030CX | 5-bromo-'2'-deoxyuridine, marker of proliferation by labelling cells during DNA synthesis |
| Mouse anti-**Ki67** | 1:500 | Abcam | Ab15580 | Marker of proliferation, labels cells active in the cell cycle |
| Rabbit anti-**S100β** | 1:500 | Dako | Z0311 | S100 calcium-binding protein β, a marker of astrocytes |
| Mouse anti-**MAP2** | 1:500 | Abcam | Ab11267 | Microtubule associated protein 2, a marker of neurons |
| Rabbit anti-**CC3** | 1:500 | Cell Signalling | 9964 | Cleaved caspase 3, marker of caspase-dependent apoptosis |
| Rabbit anti-**DCX** | 1:500 | Abcam | Ab18723 | Doublecortin, marker of neuroblasts and immature neurons |

| *Secondary Antibody* | *Dilution* | *Supplier* | *Catalogue #* |
|---|---|---|---|
| Alexa Fluor 488 Donkey anti-rat IgG | 1:500 | Life Technologies | A-21208 |
| Alexa Fluor 488 Donkey anti-mouse IgG | 1:500 | Life Technologies | A-21202 |
| Alexa Fluor 555 Donkey anti-rabbit IgG | 1:500 | Life Technologies | A-31572 |
| Alexa Fluor 555 Donkey anti-mouse IgG | 1:500 | Life Technologies | A-31570 |

**Table 1:** Top, a list of primary antibodies used in ICC experiments. Bottom, a list of secondary antibodies used in ICC experiments. For each antibody, there is also information regarding the dilution ratio used in experiments, the supplier, a catalogue number and a summary of how the marker is used (primary antibodies).

### S5: Telomere Quantification

To assess relative telomere length, we performed a modified version of a quantitative polymerase reaction (qPCR) protocol by Cawthon and colleagues, as previously described (Powell et al., 2017a; Vincent et al., 2017). The protocol involves two separate qPCRs performed on separate 384-well plates with DNA samples pipetted into identical wells on each plate. In the first reaction, we assayed the telomere repeat region (TTAGGG). In the second reaction, we assayed a single copy gene, albumin, which we used as an internal control to correct for differences in DNA concentration between samples (Cawthon, 2009). The telomere/albumin ratio was used to calculate relative telomere length.

On each plate, six negative controls consisting of RNase-free water were used to screen for any DNA contamination, and five positive controls consisting of independent leukocyte DNA samples were used to confirm a successful PCR. An eight-point dilution series using human leukocyte genomic DNA (0.47, 0.94, 1.88, 3.75, 7.5, 15, 30, and 60 ng) was used on each plate to allow for absolute quantification of each sample and to account for any differences in efficiency between the telomere and albumin reactions. All reactions were performed using three technical replicates. Each qPCR mix for the telomere reactions consisted of 10 μL of 2x qPCR Mastermix with SYBR Green (Primer Design, Southampton, United Kingdom), 5 μL or RNase free water, 12 ng of DNA, 1000 nM of telg, 5′-ACACTAAGGTTTGGGTTTGGGTTTGGGTTTGGGTTAGTGT-3′ and 800nM of telc, 5′-TGTTAGGTATCCCTATCCCTATCCCTATCCCTATCCCTAACA-3′. Four stages made up the thermocycling conditions as follows: Stage 1: 95°C for 15 min, Stage 2: 2 cycles for 15 s at 94°C and 49°C, Stage 3: 25 cycles at 94°C for 15 s, 10 s at 62°C, and 15 s at 73°C (data collection), Stage 4: dissociation curve (primer specificity detection).

The same reagents and quantities were used for the albumin reactions, apart from the albumin forward and reverse primers replaced the telomere primers. Quantities of the albumin forward and reverse primers were adjusted to 765 nM for the forward primer albu, 5′-CGGCGGCGGGCGGCGCGGGCTGGGCGGAAATGCTGCACAGAATCCTT-3′ and 930 nM for the reverse primer albd, 5′-GCCCGGCCCGCCGCGCCCGTCCCGCCGGAAAAGCATGGTCGCCTGTT-3′. The thermocycling conditions for the albumin reaction consisted of four stages: Stage 1: 95°C for 15 min, Stage 2: 2 cycles for 15 s at 94°C and 49°C, Stage 3: 33 cycles at 94°C for 15 s, 10 s at 62°C, and 15 s at 88°C (data collection), Stage 4: dissociation curve (primer specificity detection). Reactions were performed using either the QuantStudio 5 Real-Time PCR System (Thermofisher Scientific).

Quality control included checking for primer amplification specificity using melting curves, ensuring high primer efficiencies (90-110%), and confirming an $R^2$ >0.985 for all standard curves. Following this, single outliers were identified and removed if $C_t$ values corresponding to technical triplicates generated a standard deviation (SD) of above 0.5. $C_t$ values were related to absolute quantities as part of a standard curve to generate $C_q$ values. The average $C_q$ value was then generated across replicates, for each sample. To generate relative telomere length, the average $C_q$ value pertaining to the telomere repeat sequence was divided by the $C_q$ relating to the single copy gene, albumin.

### S6: hTERT Transcription levels

Telomerase reverse transcriptase (hTERT) is a catalytic subunit of the enzyme telomerase, and its expression is tightly coupled with telomerase activity (7). Prior to RNA-sequencing experiments, we checked whether there were any significant differences in *hTERT* expression between young and old cells. To do this we performed a quantitative PCR reaction using RNA samples, extracted as described in the main text.

Primers were designed systematically by exploring *hTERT* in the UCSC genome browser (https://genome.ucsc.edu), selecting an exon within *hTERT* that incorporates all transcript versions, and outputting the nucleotide sequence. This genomic sequence was then input into Primer3 (http://primer3.ut.ee; product range of between 80-150 BP), where primers were designed. Specificity of primers was confirmed *in silico*, using the ‘in silico PCR’ tool in UCSC genome browser, and validated in pilot experiments by assessing qPCR melting curves. The primers were specifically designed to generate a single amplicon, and to fall within a single exon, so that it would create the same product size for either gDNA or cDNA, allowing for the accurate use of a gDNA standard curve. Primers were supplied by Integrated DNA Technologies (IDT, London, UK).

Gene expression was normalised to both a genomic standard curve and to the reference gene Vimentin (*VIM*), as part of the relative quantification method. *VIM* is a gene that shows very low levels of variation in the hippocampal cell line as identified from microarray studies, and has previously been used as a reference gene in cells grown across multiple passages (2).

1 µg of total RNA from each sample was submitted to genomic DNA wipeout using DNA-free™ DNA Removal Kit (Invitrogen, California, USA). This was carried out to increase the purity of RNA, prevent non-specific binding of primers and achieve optimal qPCR results. gDNA-free RNA was the submitted to reverse transcription using the Invitrogen SuperScript III Reverse Transcriptase kit (Invitrogen, California, USA), to generate complementary DNA (cDNA). cDNA samples were then diluted to a working concentration of 5ng/µl.

qPCRs were performed on 384-well plates with each sample plated in triplicate. Per gene, we used an eight-point standard curve consisting of DNA samples at 0.47 ng, 0.94 ng, 1.88 ng, 3.75 ng, 7.5 ng, 15 ng, 30 ng, 60 ng amounts. This allowed us to correct for any minor differences in PCR efficiency between our target (*hTERT*) and reference gene (*VIM*); though all PCR efficiencies were between 90-110 %. We also included a no-template control consisting of RNase free water to check for nucleic acid contamination.

The final 12.5 µl qPCR reaction consisted of 25 ng of cDNA – 5 µl, 1x EvaGreen – 2.5 µl of 5x mix, 200 nM of forward and reverse primers – 0.25 µl, and 4.75 µl of water, to make up 12.5 µl total volume. Primer sequences for the *hTERT* reaction consisted of forward, 5' - CAGGATGGAGTAGCAGAGGG - 3', and reverse primers, 5' - GCAGCTCCCATTTCATCAGC - 3'. Primer sequences for the *VIM* reaction consisted of forward, 5' - CTTTGCCGTTGAAGCTGCTA - 3', and reverse primers, 5' - ACGAGCCATTTCCTCCTTCA - 3'. The thermocycling conditions were as follows: 15 min at 95 °C (denaturation and Taq enzyme activation), then 40 cycles of: 30 sec at 95 °C, 30 sec at 60 °C, and 30 sec at 72 °C (data collection), followed by a melting curve. qPCRs were run on the QuantStudio 5 Realtime PCR system (Thermo Fisher Scientific).

Cycle threshold ($C_t$) values were the primary data produced, and correspond to how many PCR cycles were required for a sample to reach a predefined level of fluorescence. Single outliers from triplicates were identified and removed, where SD > 0.5. $C_t$ values were then related to absolute quantities as part of a standard curve to generate $C_q$ values. The average $C_q$ value was then generated across replicates, for each sample. For relative gene expression analyses the average $C_q$ pertaining to the target gene, *hTERT*, was divided by the $C_q$ relating to the reference gene, *VIM*.

Finally, a two-tailed t-test was used to draw comparisons between old and young cells, which revealed no significant differences in *hTERT* expression between young and old cells ($P > 0.05$).

**S7: Telomere length changes in the fetal brain**

To confirm whether our cell ageing model in neural progenitors could also be capturing a neurodevelopment process, we tested whether there was a relationship between post-conception days (a proxy for cell division number) and telomere length, in foetal brain samples.

Human foetal brain samples were provided by the Human Developmental Biology Resource (HDBR) (http://www.hdbr.org). Ethical approval for the HDBR was granted by the Royal Free Hospital research ethics committee under reference 18/NE/0290. Brain tissue was obtained frozen and had not been dissected into regions. Half of the brain tissue from each individual foetus was homogenised for subsequent genomic DNA extraction, which was performed by standard phenol-chloroform procedures as described previously (8). Sample sex was determined via PCR amplification (8). In total, DNA was available from 37 females and 48 males (n=85). Gestational ages were estimated using foot and knee to heel length measurements. Gestational ages ranged from 75-161 days, with a mean gestational age of 107.96 (S.D. = 16.12). Telomere length was quantified as described in S5.

To test the effect of gestational age on telomere length in the fetal brain, we ran a linear regression with relative telomere length as the outcome variable, sex as a covariate, and gestational age (days) as the independent variable. We found no significant relationship between gestational age and telomere length in fetal brain ($\beta$ = -0.003 [95% CI: -0.328 – 0.322], P = 0.985), Figure 1. We also found no sex differences in telomere length from the fetal brain ($\beta$ = -0.001 [-0.011 – 0.009], p = 0.887). This suggests that telomere shortening observed in the hippocampus is unlikely to relate to a prenatal neurodevelopmental processes.

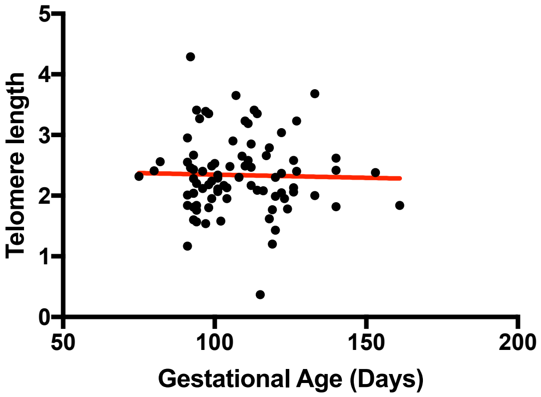


Figure 1: Gestational age (a proxy for number of cell divisions) does not significantly correlate with telomere length in the foetal brain.

<u>References</u>

1. Powell TR, *et al.* (2017) Transcriptomic profiling of human hippocampal progenitor cells treated with antidepressants and its application in drug repositioning. *J Psychopharmacol* 31(3):338-345.
2. Powell TR, *et al.* (2017) Inter-individual variation in genes governing human hippocampal progenitor differentiation in vitro is associated with hippocampal volume in adulthood. *Scientific Reports* 7(1):15112.

3. Powell TR, *et al.* (2017) The genome-wide expression effects of escitalopram and its relationship to neurogenesis, hippocampal volume, and antidepressant response. *Am J Med Genet B Neuropsychiatr Genet* 174(4):427-434.
4. Smeeth DM, Kourouzidou I, Duarte RRR, Powell TR, & Thuret S (2020) Prolactin, Estradiol and Testosterone Differentially Impact Human Hippocampal Neurogenesis in an In Vitro Model. *Neuroscience*.
5. Anacker C, *et al.* (2011) Antidepressants increase human hippocampal neurogenesis by activating the glucocorticoid receptor. *Mol Psychiatry* 16(7):738-750.
6. Wang J, Xie LY, Allan S, Beach D, & Hannon GJ (1998) Myc activates telomerase. *Genes Dev* 12(12):1769-1774.
7. Leao R, *et al.* (2018) Mechanisms of human telomerase reverse transcriptase (hTERT) regulation: clinical impacts in cancer. *J Biomed Sci* 25(1):22.
8. Spiers H, *et al.* (2015) Methylomic trajectories across human fetal brain development. *Genome Res* 25(3):338-352.
